# The Ras family members follow the blood progesterone level during formation and regression in bovine corpus luteum

**DOI:** 10.1101/625186

**Authors:** Sang-Hee Lee, Seunghyung Lee

**Author notes:** This document certifies that the manuscript listed below was edited for proper English language, grammar, punctuation, spelling, and overall style by one or more of the highly qualified native English speaking editors at American Journal Experts (Verification Key No: 4A04-0DC7-76BA-A4B0-BEC9).

## Abstract

Ras family members regulate cellular differentiation, proliferation and survival. CL formation and regression are regulated by the blood P4 level. This study investigated the association between changes in Ras family members and the serum P4 level and determined protein interactions among Ras family members, hormone receptors, and angiogenetic and apoptotic factors during formation and regression of the bovine CL. RASAL3 and RASA3 were found using proteomics in CL and were significantly increased in the SPCL compared to the PPCL, whereas RasGEF1B was decreased in the PPCL. Hormone receptors and angiogenetic proteins expression was lower in the PPCL and SPCL than that in the RPCL, but apoptotic proteins were increased in the RPCL. The P4 and estrogen receptors positive correlated with RasGEF1B, R-Ras, and H-Ras through VEGFA, VEGFR2 and Tie2 in STRING database. RasGAP, H-Ras and R-Ras protein expression was increased in the PPCL compared to that in the SPCL, whereas RasGEF expression was decreased. In summary, Ras activation and angiogenesis in the CL were positively correlated with the blood P4 during estrous cylce. These results may increase understanding of Ras biological functions following stimulation of hormones and their receptors during tissue proliferation and degeneration.

## Introduction

The corpus luteum (CL) is a transient endocrine gland in the female reproductive tract that produces progesterone (P4), which is required to maintain pregnancy for the beginning of life in mammals (1). During CL formation, granulosa and theca cells in the ovary are differentiated and proliferate into steroidogenic luteal cells (LSCs) in response to luteinizing hormone (LH) until day 8 to 9 after ovulation. Then, the CL weight increases three to four times when its growth is complete prior to the next round of ovulation (2). LSCs produce P4 that is secreted into the blood vessels, which contribute to the maintenance of pregnancy (3–5). However, if the pregnancy is not established, the CL begins to regress in response to prostaglandin F2 alpha (PGF2α) derived from the endometrium in a process termed luteolysis (6). The formation and regression of the CL are repeated, and this repetition by sex hormones distinguishes it from other mammalian tissues (6). For several decades, studies have mainly investigated the formation and regression of gonadotropin release hormone (GnRH) and steroidogenic hormones, whereas few studies have investigated the roles of small GTPases in formation and regression of the CL.

During the proliferation phase (PP), luteal endothelial cells (LECs) proliferate following interaction with vascular endothelial growth factor A (VEGFA) and VEGF receptor 2 (VEGFR2) to generate blood vessels and support LSC proliferation in the CL (7, 8). After vasculature is stabilized by the balance of angiopoietin 1 (Ang1) and Tie2 (9). When the CL is completely developed, P4 is converted from pregneolone by 3β-hydroxysteroid dehydrogenase (3β-HSD) continuously during a phase that is called the secretion phase (SP) (6). If implantation is successful, maternal recognition should inhibit PGF2α synthesis in the endometrium, and then the CL consistently produces and secretes P4 to maintain the pregnancy (10). However, synthesized PGF2α from the endometrium is transported through blood vessels and triggers CL regression when the endometrium does not recognize maternal processes; thus, the purpose of luteolysis induced by PGF2α is to prepare for a new estrous cycle (3, 11). During luteolysis as a regression phase (RP), representative cell death signals, tumor necrosis factor (TNF), and FasL and its receptors are activated. The P4 concentration dramatically increases, and the PGF2α concentration increases in the blood and CL (12). Therefore, the blood P4 and PGF2α concentrations are key points for the proliferation, angiogenesis, and apoptosis of luteal cells.

Small G proteins are typically between 20 and 30 kDa in size and cycle between an inactive guanosine diphosphate (GDP)-bound conformation and an active guanosine triphosphate (GTP)-bound conformation that act as molecular switches to regulate broad cellular processes, including proliferation, differentiation, adhesion, survival, and apoptosis (13). The GDP- and GTP-conformation cycle is regulated by guanine nucleotide exchange factors (GEFs), which induce the release of bound GDP and its replacement with GTP by GTPase-activating proteins (GAPs) that provide a catalytic group for GTP hydrolysis (13). The progenitor of the small G-protein family is Ras, which is mutated in 15% of human tumors. Ras consists of over 150 members that are classified into families and subfamilies based on sequence and functional similarities. The Ras superfamily is classified into five principal families (Ras, Rho, Rab, Arf, and Ran) (14). The Ras family members regulate various signaling pathways, including those involved in transcription, cellular differentiation and proliferation (13).

Although LSCs and LECs continually differentiate and proliferate and P4 gradually increases after ovulation during CL formation (7, 8), few studies has investigated the effect of Ras family members on differentiation and proliferation or the relationship with the P4 concentration in CL during the estrous cycle (15, 16). Therefore, this study investigated the blood P4 level and discovered Ras regulator proteins during the estrous cycle in the bovine CL based on protein correlation. Then, to evaluate the association between Ras regulators and hormone receptors, protein-protein interactions among the discovered Ras regulators, hormone receptors, angiogenetic factors and apoptotic factors were analyzed using bioinformatics methods. Lastly, correlative relationships of mediator factors between Ras family members and hormone receptors were investigated, and their activation was comparatively analyzed with changes in the P4 concentration and tissue formation and regression during the estrous cycle.

## Materials and methods

### Animals, progesterone levels in blood and sample preparation

All procedures involving animals were approved by the Kangwon National University Institutional Animal Care and Use Committee (KIACUC-09-0139). Estrous synchronization and detection were performed as described previously (17). To induce estrous synchronization, exogenous progesterone source, Controlled Internal Drug Released dispenser (CIDR; Pfizer Animal Health, New York, NY, USA), was inserted in vagina, and GnRH (Pfizer) injected in cows (*n* = 10). After 48 hours, CIDR was removed to regress P4 in blood, then PGF2α was injected for CL regression and new estrous cycle. Estrous sign was observed at 36-48 hours after PGF2α injection, this point was defined as the ovulation (Day 0). Blood was collected from jugular venipuncture using 15 mL vacutainer cleaned by heparin, between 10:00 to 11:00 am at every 2 days from Day-2 to 24. Collected blood were centrifuged at 3,000 rpm for 15 min at 4°C, and serum was isolated and stored in −80 °C until the analysis. P4 levels were measured by ESLIA kit (Enzo Life Sciences, New York, NY, USA) according to the manufacturer’s instructions. CL were collected from slaughtered heifers, transferred in laboratory within 2 hours at 4°C. Collected CL samples were classified according to three morphology such as 1-2 days (proliferation phase CL; PPCL; *n* = 15), 12-15 days (secretion phase CL; SPCL; *n* = 20) and 18-20 days (regression phase CL; RPCL; *n* = 23) after ovulation. Then, isolated CL tissues from ovary were measured weight and stored at −80°C until experiment.

### Hematoxylin and eosin staining

Fixation and paraffin section of CL tissues were conducted according to previously methods (18). The fixed and paraffin embedded CL tissues were 4 μm sections using a microtome. The paraffin sections were deparaffinized in xylene and rehydrated in ethanol (100%, 90%, 80%, and 70%) for 5 min. The CL tissue was blocked by immersing the slides in 3% BSA in PBS for 60 min and then washed with PBS for 5 min at room temperature (RT), then samples were stained in hematoxylin solution (Sigma, St Louis, MO, USA) for 5 min and washed with distilled water for 10 min, stained in EosinY solution (Sigma) for 1 min. Then samples were washed with distilled water, and dehydrated in ethanol (70%, 80%, 90% and 100%) and 100% xylene for 5 min per each step. The slides were mounted with Histomount Mounting Solution (Thermo Scientific, Waltham, MA, USA), then observed using Olympus BX50 microscope (Olympus, Tokyo, Japan).

### Two-dimensional gel electrophoresis (2-DE)

CL samples (PPCL, *n* = 4; SPCL, *n* =4; RPCL, *n* =4) were selected from isolated CL tissues from ovaries according *3β-HSD* mRNA expression (Supplementary S1 Fig). 3β-HSD expression means P4 production from CL because P4 is converted from pregneolone by 3β-HSD (1). CL samples were homogenized in M-PER mammalian Protein Extraction Reagent (Thermo Scientific) using tissue homogenizer (Bioneer, Daejeon, Korea), then incubated for 1 hour at RT, and centrifuged at 12,000 g for 10 min at 4 °C. The supernatant was transferred into a new micro-centrifuge tube, and protein concentration was determined by the BCA protein assay kit (Thermo Scientific). Interfered with substances such as detergents, slats, lipids, phenolics and nucleic acids in extracted protein were removed using the 2-D Clean-Up kit (GE Healthcare, Piscataway, NJ, USA) according to the manufacturer’s instructions, and 700 μg protein was dissolved in 300 μL rehydration solution (GE Healthcare) for 1 hour at RT. Proteins into rehydration buffer were incubated with an 18 cm immobilized pH 3–11 nonlinear gradient dry strip (GE Healthcare) for 16 hours at 20°C. As described previously (19), isoelectric focusing (IEF) was performed for protein separation. IEF was performed at hold 500 V for 1 hour, gradient 1,000V 1hour, gradient 8,000 V for 3 hours, hold 8,000 V for 1.5 hours, gradient 10,000 V for 3 hours and hold 10,000 V for 1hour. Strips were then equilibrated for 15 min in 5 mL equilibration buffer (50 mM Tris-HCl, pH 8.8, 6.0 M urea, 30% glycerol (v/v), and 2% sodium dodecyl sulfate (w/v) containing 0.8 g dithiothreitol (Sigma), followed by an additional incubation for 15 min in 5 mL equilibration buffer containing 0.1 g iodoacetamide. Separation in the second dimension was accomplished using an 10% SDS-PAGE in a Protean II xi 2-D Cell (Bio-Rad, Hercules, CA, USA) at 50 mA until the bromophenol blue reached the bottom of the gel. Gels were stained in a solution of 0.1% Coomassie Brilliant Blue R-250 (Sigma) comprised of 45% methanol, 10% acetic acid and 45% water. Gels were then scanned using an image scanner and analyzed with ImageJ software (NCBI, USA). The SPCL and RPCL protein spots intensities were normalized to PPCL protein spots intensities for calculation of relative intensity.

### Matrix-assisted laser desorption/ionization time-of-flight mass spectrometry (MALDI-TOF/MS)

MALDI-TOF/MS was performed as described previously (19) and condition was listed in Supplementary Table 1. Spots were extracted from the gel and washed in 50% acetonitrile (ACN; Sigma) containing 25 mM NH_4_ bicarbonate, then incubated with 50% ACN containing 10 mM NH_4_ bicarbonate and 100% CAN. Finally, ACN in samples was removed using a speed vacuum. Then, samples were incubated with cold sequencing-grade modified trypsin (Promega, Madison, WI, USA) at 37°C for 20 h, followed by 50 min incubation with 50% ACN containing 5% trifluoroacetic acid (TFA) at RT. The supernatants were dried for peptide extraction using a speed vacuum, and then diluted with 50% ACN containing 5% TFA. Samples were desalted using a Zip-Tip C18 (Millipore, Milford, MA, USA). Plating was performed using a 4-hydroxy-α-cyano-cinnamic acid matrix solution (Sigma) on a MALDI-TOF/MS plate. Peptides were analyzed using an Ultraflex-TOF/TOF spectrometer (Bruker Daltonics, Hamburg, Germany) and MS-Fit software (http://prospector.ucsf.edu) and data were searched against UniProt database (http://www.uniprot.org/).

### Bioinformatics analysis

Ras regulator proteins of CL were analyzed to detection protein-protein interaction with **i**) hormone receptors such as P4 receptor (P4R), PGF2α receptor (PGF2αR), estrogen receptor alpha (ERα), and oxytocin receptor (OTR), **ii**) angiogenesis factors such as VEGFA, VEGFR2, Ang1, Tie 2 and hypoxia inducible factor 1 alpha (HIF1α), **iii**) apoptosis factors such as TNF receptor 1 (TNFR1), Fas, Bax, Bcl-2, caspase 3 (casp3) and p53, and **iV**) Ras proteins such as transforming protein p21 (H-Ras) and Ras-related protein (R-Ras). A Protein-protein interaction analysis of the identified proteins was performed using Search Tool for the Retrieval of Interacting Genes (STRING) database v.10.0 (http://string.embl.de) with the flowing analysis parameters Bos taurus species and all interaction sources. Biological processes, molecular function, and cellular components of CL proteins were classified as Gene Ontology (GO) of STRING functional enrichment network. The PPCL, SPCL and RPCL were preferentially analyzed using network and molecular action tools, then classified as Ras regulators according to biological process and molecular function in the STRING enrichment network system. Protein-protein interaction were shown using confidence value that is complex of the textmining, experiments, databases, co-expression, neighborhood, gene fusion and co-occurrence based on STRING database.

### Quantitative RT-PCR

The mRNA was extracted using TRIzol (Takara, Shiga, Japan), and concentration was measured using NanoDrop 2000 spectrophotometer (Thermo Scientific). Total 5.0 μg mRNA was used to synthesis cDNA using PrimScript 1^st^ strand cDNA synthesis kit (Takara), and reverse transcription was performed at 45°C for 60 min after 95°C for 5 min. The 1.0 μL synthesized cDNA were used to conduct PCR and was performed according to the primer conditions (Supplementary Table S2) using PCR premix kit (Bioneer). Then, the products were separated with 2.0% agarose gel electrophoresis at 100 V for 20 min, stained with ethidium bromide, visualized with UV light, and mRNA expression was analyzed with ImageJ software (NCBI).

### Western blot

The proteins (25 μg/20 μL) were separated by sodium dodecyl sulfate–polyacrylamide gel electrophoresis (SDS-PAGE) at 30 V for 20 min after 100V for 90 min, transferred to a polyvinylidene difluoride (PVDF) membrane at 30 V for 180 min at 4 °C, and incubated in blocking solution (5% skim milk in Tris-buffered saline/0.5% Tween-20; TBS-T) at RT for 60 min. The membranes were incubated with TBS-T with 1 % bovine serum albumin (BSA; Sigma) containing primary antibodies at 4 °C for overnight. The membranes were then washed three times with TBS-T each 5 min and incubated with secondary antibodies conjugated horseradish peroxidase and visualized using the West Save Enhanced Chemiluminescence kit (AbFrontier, Austin, TX). Protein expression was measured using the EZ-Capture II system (ATTO, Tokyo, Japan), and protein band intensity was calculated using ImageJ software (NCBI). Used antibodies were listed in Supplementary Table S3.

### Statistical analyses

Data were analyzed using SAS ver. 9.4 software (SAS Institute, Cary, NC). Data are presented as mean ± standard error. Data were evaluated using analysis of variance (ANOVA) and Duncan’s multiple-range test using general linear models. A P value < 0.05 was considered to indicate statistical significance.

## Results

### 2-DE, mass spectrometry, and protein association analysis

Total 56 protein spots were detected in CL (Supplementary Table S4), of these 27 protein spots were repetitively detected at PPCL (Fig 1B), SPCL (Fig 1C) and RPCL (Fig 1*D*) were analyzed by MALDI-TOF/MS (Supplementary Fig S2; spot no. 1-12 and S3 Fig; spot no. 13-27) and determined to correspond to 18 different proteins (Table 1) which processing is summarized in Fig 1*A*. The confidence value of RABL5, RASAL3, RASA3, RasGEF1B, GSTA1, SWAP70, and GDI2 were less with other 11 proteins (Fig 1E). The RABL5 is play a role intracellular protein transport and GTP binding, RASAL3 and RASA3 (Table 1). The RASAL3, RASA3, RasGEF1B, and GDI2 play a role positive regulation of GTPase activity (Fig 1F, red), RABL5, RasGEF1B, and GDI2 mediate signal transduction vial small GTPase (Fig 1F, blue), and RASAL3, RASA3 and GDI2 active GTPase activators (Fig 1F, green). Especially, RASAL3, RASA3, and RasGEF1B were directly involved in Ras proteins activation compared to RABL5 and GDI2 (Table 1). Based on 2-DE protein spot analysis, RABL5, RASAL3, RASA3, RasGEF1B and GDI2 protein spots were decreased in RPCL compare to PPCL and SPCL (Fig 1G). Especially, RASAL3 and RASA3 were increased but RasGEF1B were decreased in SPCL compared to PPCL (Fig 1*G*).

**Fig 1.**
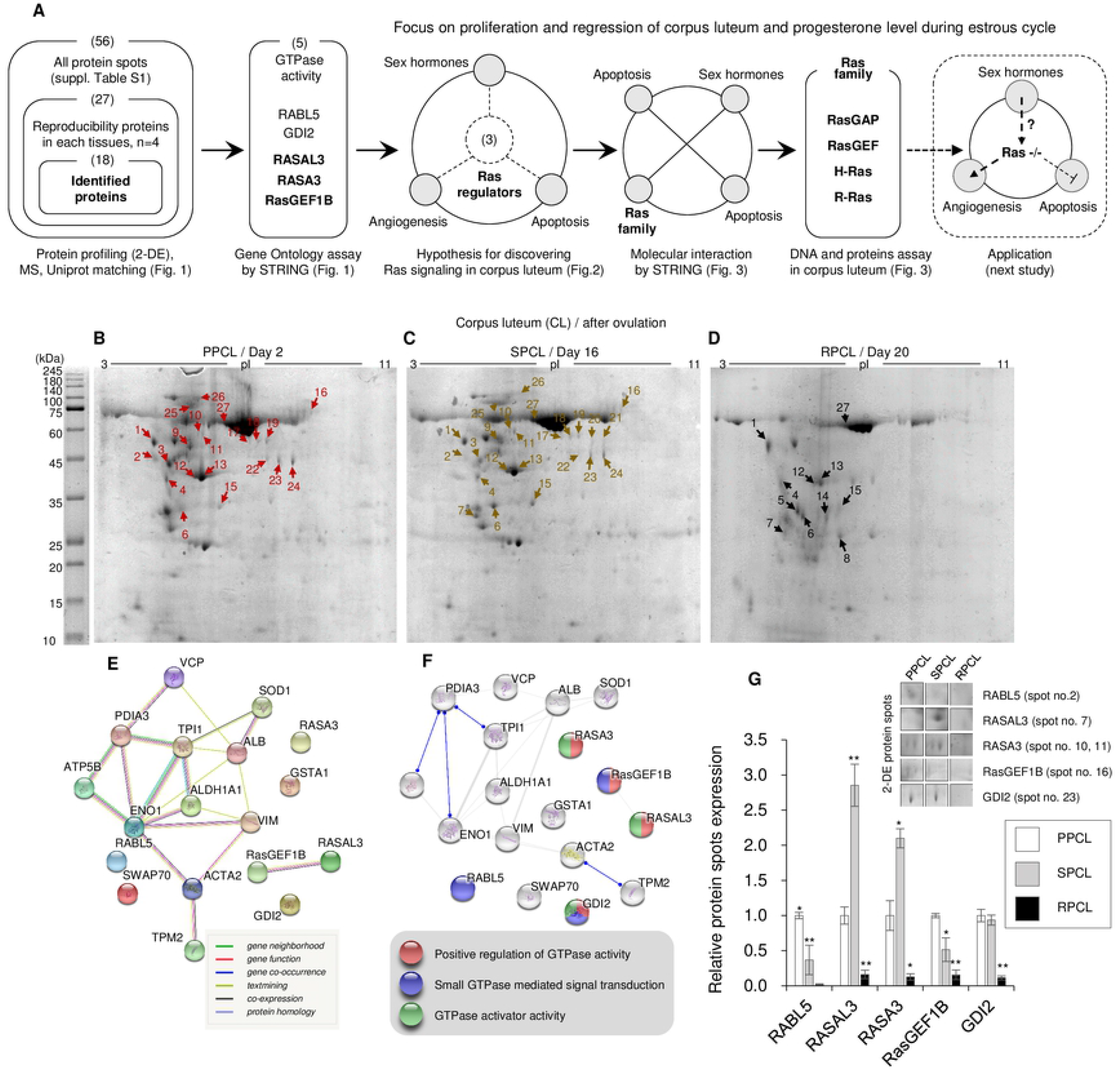
Strategy of discover on Ras family members signaling focused on proliferation and regression of corpus luteum (CL) and progesterone level during estrous cycle based on two dimensional electrophoresis (2-DE), mass spectrometry (MS), and bioinformatics (A), Distribution of protein spots in acrylamide gels using 2-DE of the CL at day 2 (B, proliferation phase CL; PPCL), day 16 (C, secretion phase SPCL; SP), and day 20 (D, regression phase CL; RPCL) after ovulation (each phase CL, *n* = 4). The spot numbers correspond to the labels in Table 1. The original 2-DE gel image with size marker were shown as Supplementary Fig S9. The protein network of the discovered protein spots was analyzed using evidence tools in the STRING database (E, F). The line color indicates the type of interaction evidence. (F) Molecular interaction of protein spots in the CL during the estrous cycle. The blue line indicates the binding ability, and the line shape indicates the predicted mode of action. Proteins related to GTPase activity were classified with the gene ontology (GO) tool in the STRING database. (G) Changes in RABL5 (Spot no. 2), RASAL3 (Spot no. 7), RASA3 (Spots no. 10 and 11), RasGEF1B (Spot no. 16), and GDI2 (Spot no. 23) protein spot expression at the PPCL, SPCL, and RPCL. Expression of protein spots at the SP and RP was normalized to that of the PP. **p<0.01 and *p<0.05, n=4.

**Table 1.**
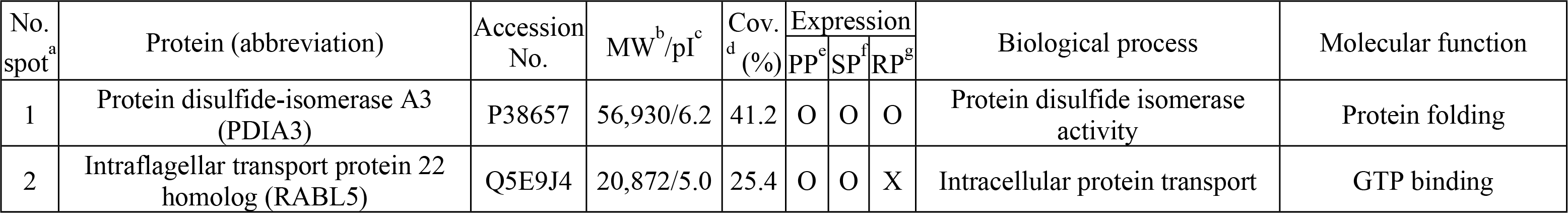
Change of proteins among corpus luteum during estrous cycle

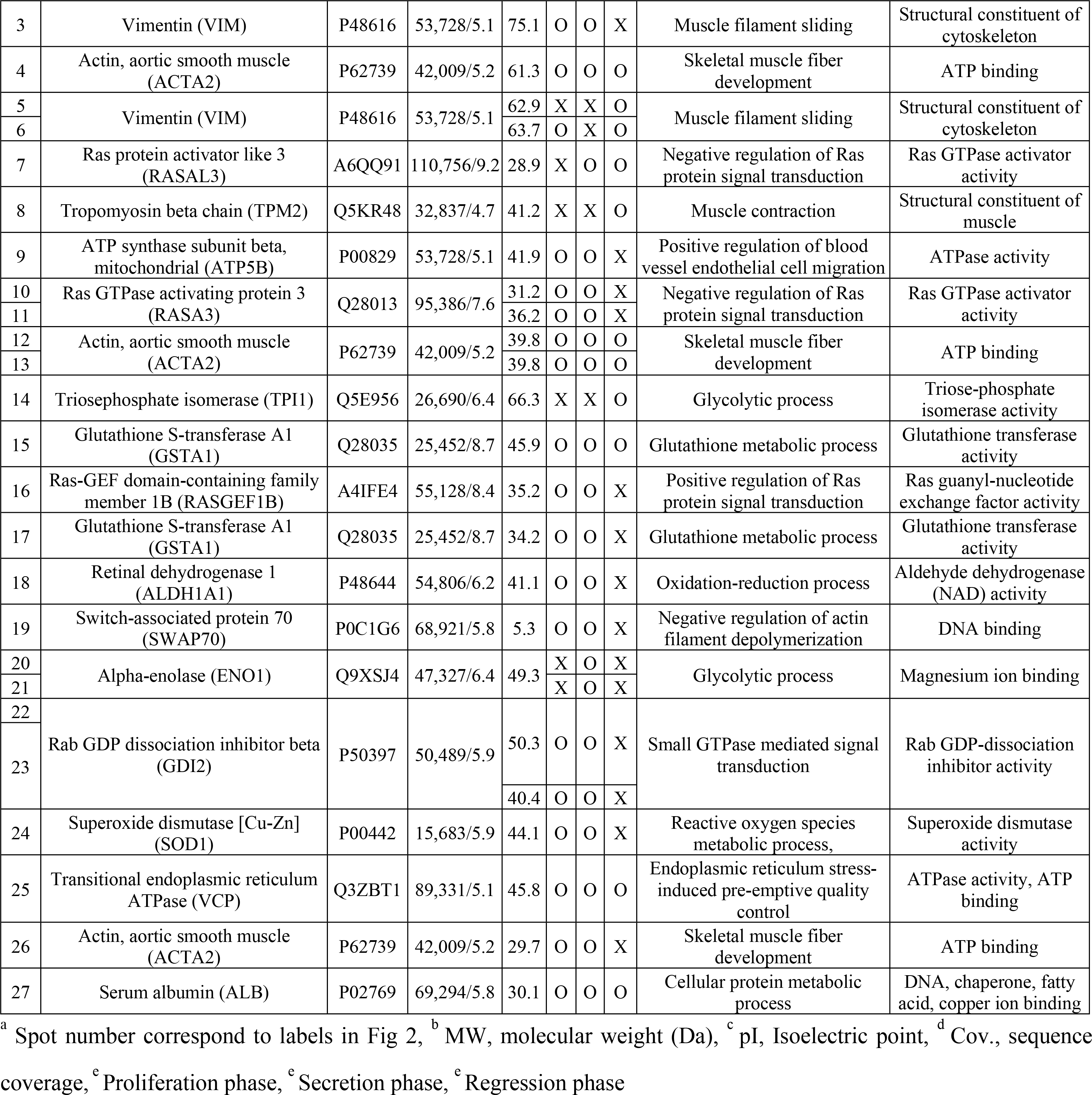

The Gene Ontology (GO) analysis of identified total CL proteins reveals over the half of all proteins at PPCL and SPCL involved in cellular process, single organism cellular process, biological process, cellular response to stimulus, and cell communication, whereas apoptotic process, negative cellular process, negative biological process, endocytosis, negative developmental process, and muscle contraction were mostly related with RPCL (Fig 2A). Based on classification of the identified differentially expressed protein in molecular function, over the 50 % of PPCL and SPCL proteins were involved in molecular function, heterocyclic compound binding, organic cyclic compound binding and ion binding (Fig 2B). Moreover, cellular components ratio of determined proteins was different during estrous cycle, kinds of cellular component were increased in PPCL (Fig 2C) to SPCL (Fig 2D), whereas, decreased at SPCL to RPCL (Fig 2E).

**Fig 2.**
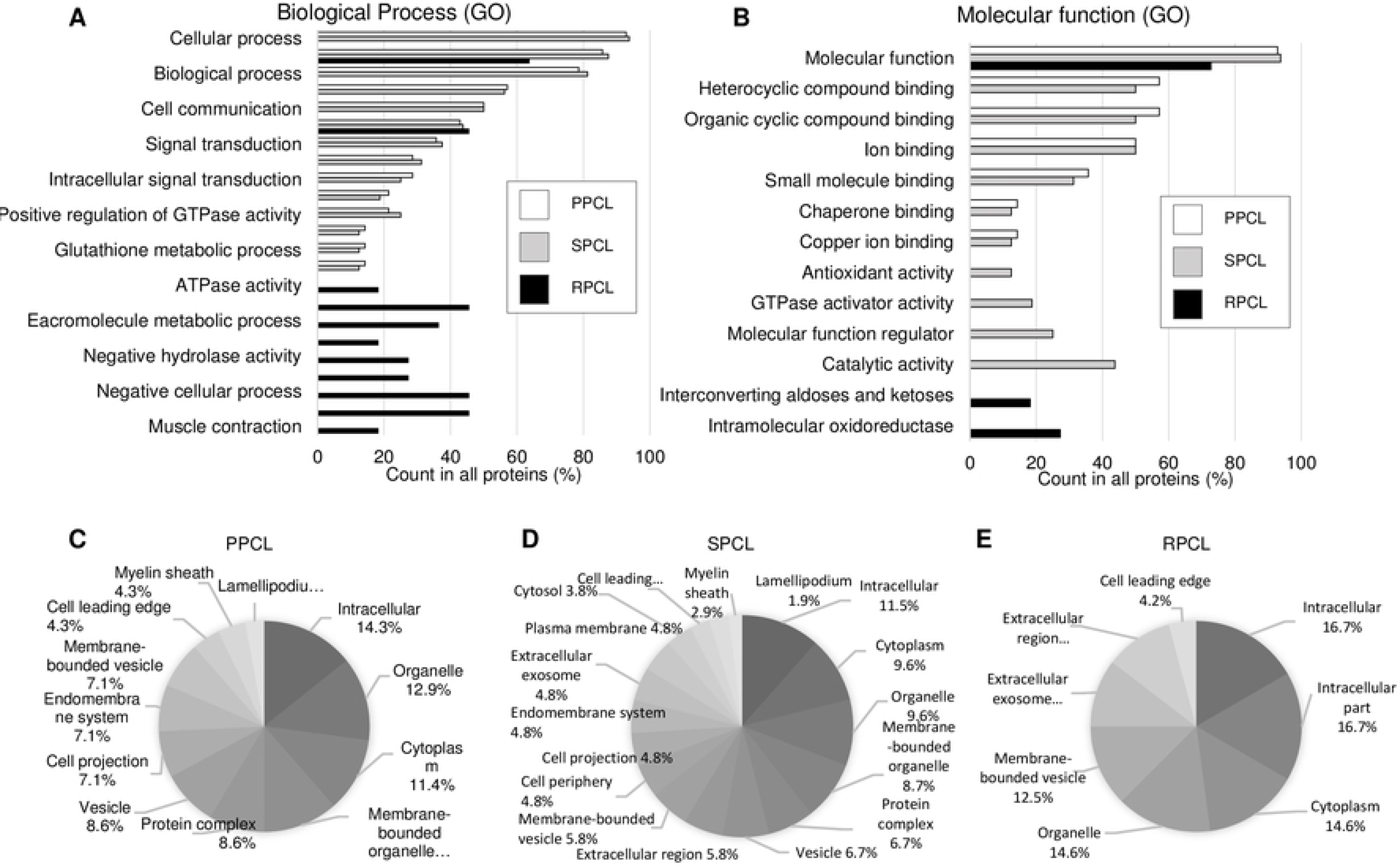
Biological processes (A), molecular functions (B) and cellular component ratios (C) of corpus luteum (CL) protein spots at proliferation phase (PP) CL, secretion phase (SP) CL, and regression phase (RP) CL were analyzed using Gene Ontology, and (C-E) the cellular component ratios of the protein spots from the PPCL (C), SPCL (D), and RPCL (E) were analyzed using the STRING database.

### Protein connection in hormone receptors, angiogenesis, apoptosis factors and Ras regulators

The 3β-HSD, P4R, PGF2αR and ERα mRNA (Fig 3A) and protein (Fig 3B) were decreased in RPCL compared to PPCL and SPCL, ERα and OTR proteins expression were higher at PPCL than SPCL and RPCL. Additionally, P4R was positive correlated with ERα but there were little molecular interaction between hormone receptors and Ras regulators according to in STING protein-protein interaction data (Fig 3C).

**Fig 3.**
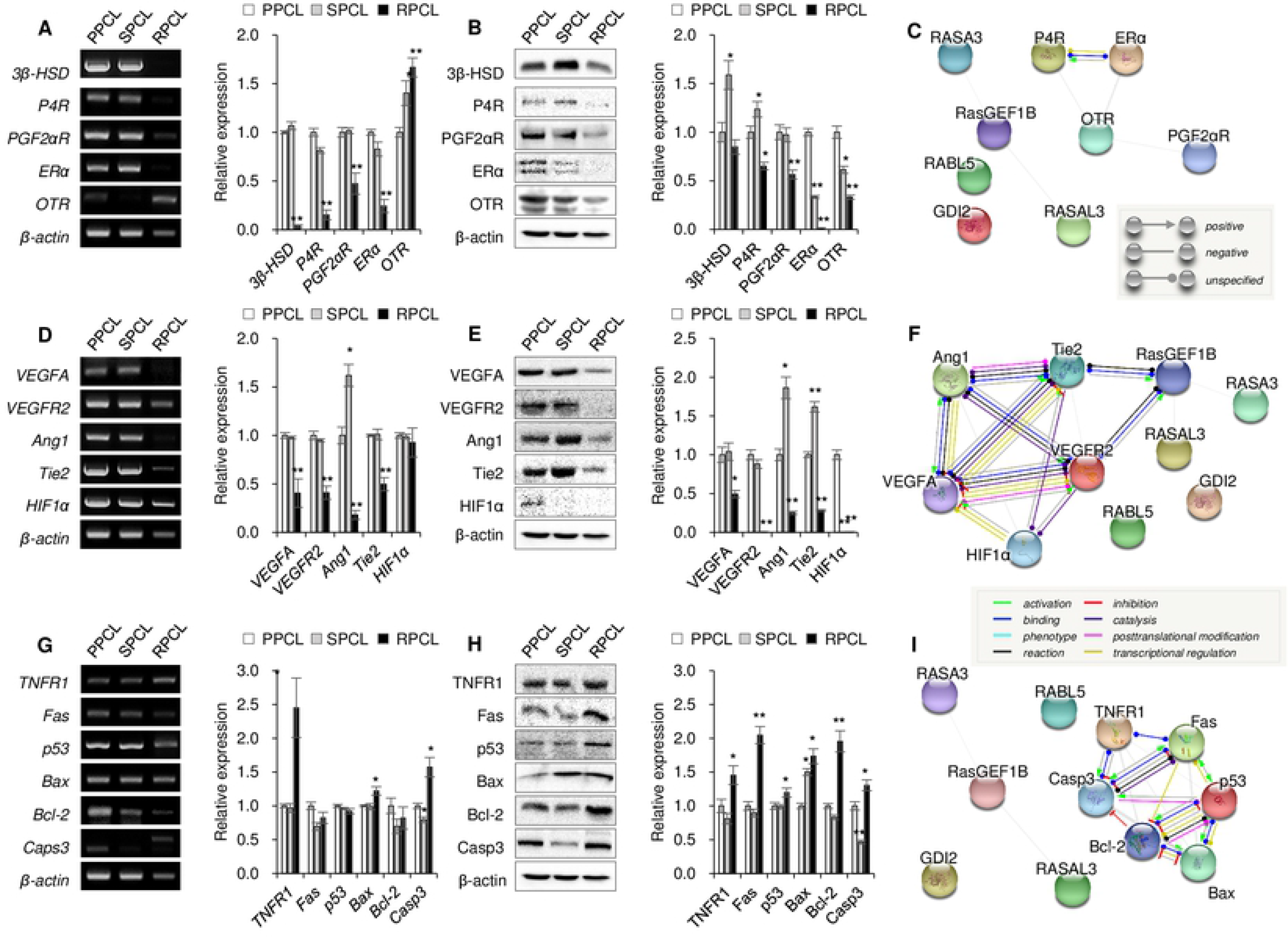
Association of hormone receptors, angiogenesis, apoptosis and Ras family member proteins in corpus luteum during estrous cycle. Changes in 3β-Hydroxysteroid dehydrogenase (3β-HSD), progesterone receptor (P4R), prostaglandin F2 alpha receptor (PGF2αR), estrogen receptor alpha (ERα) and oxytocin receptor (OTR) mRNA (A) and protein (B); vascular endothelial growth factor A (VEGFA), VEGF receptor 2 (VEGFR2), angiopoietin 1 (Ang1), Tie2, and hypoxia inducible factor 1 alpha (HIF1α) mRNA (D) and protein (E); tumor necrosis factor receptor 1 (TNFR1), Fas, p53, Bax, Bcl-2 and caspase 3 (Casp3) mRNA (G) and protein (H) expression at the proliferation phase (PP, *n* = 4), secretion phase (SP, *n* = 4), and regression phase (RP, *n* = 4) during the estrous cycle in the corpus luteum (CL). mRNA and protein expression at the SPCL and RPCL, was normalized to that of the PPCL. Molecular action between GTPase regulators (RABL5, RASAL3, RASA3, RasGEF1B, and GDI2) and hormone receptors (C; P4R, PGF2αR, ERα, and OTR), angiogenetic factors (F; VEGFA, VEGFR2, Ang1, Tie2, and HIF1α) and apoptotic factors (I; TNFR1, Fas, p53, Bax, Bcl-2 and Casp3). The line shape indicates the predicted mode of action. **p<0.01 and *p<0.05. The size and molecular weight image of PCR products and proteins were shown as Supplementary Fig S10-S13.

The VEGFA and VEGFR2 mRNA (Fig 3D) and protein (Fig 3E) were increased at PPCL and SPCL compared to RPCL. The Ang1 mRNA and protein expression were higher at SPCL than PPCL and RPCL, and Tie2 mRNA was no significantly difference between PPCL and SPCL, but protein was increased at SPCL compared to PPCL and RPCL. (Fig 3E). According to STING data, there were highly correlative between angiogenetic factors and Ras regulators, moreover both VEGFA and Tie2 had various molecular action to VEGFR2 and Tie2 (Fig 3F). Furthermore, STRING data showed that the VEGFR2 and Tie2 were not only bound (blue line) and reacted (black line) with RasGEF1B but also was activated (green arrow) by the VEGFR2 and Tie2 (Fig 3F).

The TNFR1, Bax and Casp3 mRNA (Fig 3G) and protein (Fig 3H) were increased at RPCL than PPCL and SPCL. Fas protein were increased at RPCL, but mRNA were not different among the CL phases. There were few molecular action between apoptotic factors and Ras regulators in STRING database (Fig 3I).

### Association of hormone receptors, angiogenesis, apoptosis and Ras family member proteins in corpus luteum during estrous cycle

Protein association among hormone receptors, angiogenetic factors, apoptotic factors, Ras regulators (RASAL3, RASA3 and RasGEF1B), and Ras proteins (H-Ras and R-Ras) were shown Fig 4A. According to STRING database, P4R and ERα were activated (green arrows) with p53 each other, and ERα had transcriptional regulation (yellow line) to VEGFA, and the RasGEF1B were activated, bound (blue line) and reacted (black line) by VEGFR2 and Tie2 (Fig 4A). Additionally, H-Ras and R-Ras were catalyzed (purple line), bound and activated by VEGF2, Tie2 and RasGEF1B, H-Ras had many molecular action with VEGFA distinct from R-Ras (Fig 4A) according to STRING database. Interestingly, only H-Ras of the Ras family member was bound and activated by p53 and inhibited (red line) Casp3 in apoptotic factors (Fig 4A). RasGAP, RasGEF and Ras mRNA (Fig 4*B*) and proteins (Fig 4C) were decreased in RPCL compared to PPCL and SPCL. The RASA3, RASAL3, and RasGEF1B mRNA were no significantly between PPCL and SPCL, H-Ras and R-Ras mRNA expression were reduced in SPCL and RPCL compared to PPCL (Fig 4B). The RasGAP protein expression were higher in SPCL than PPCL, otherwise RasGEF, H-Ras, and R-Ras were reduced in SPCL and RPCL compared PPCL (Fig 4C).

**Fig 4.**
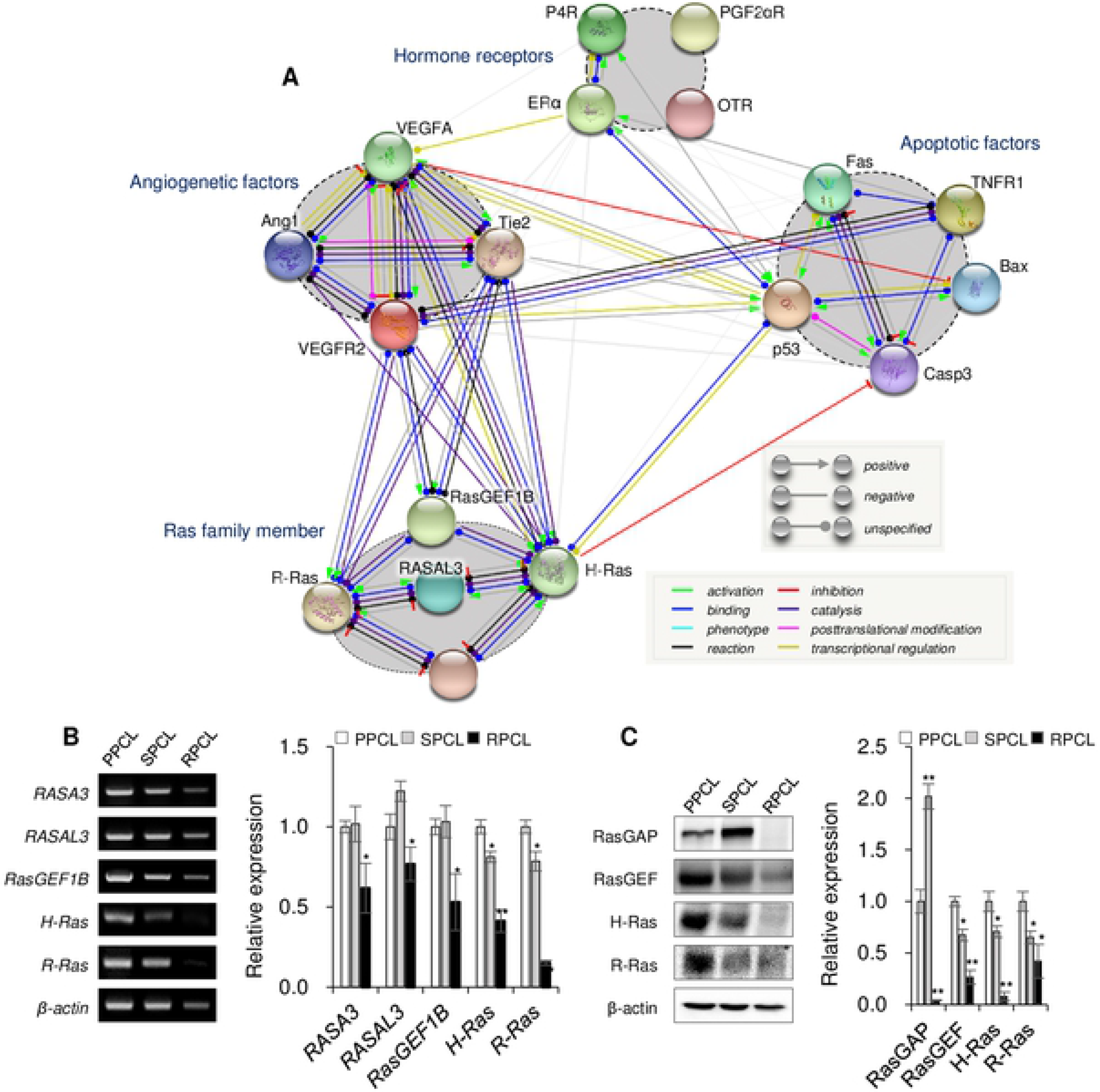
Association of hormone receptors, angiogenesis, apoptosis and Ras family member proteins in corpus luteum during estrous cycle (A). Molecular action among hormone receptors (P4R, PGF2αR, ERα, and OTR), angiogenetic factors (VEGFA, VEGFR2, Ang1 and Tie2), apoptotic factors (TNFR1, Fas, p53, Bax, and Casp3), Ras regulators (RASAL3, RASA3, and RasGEF1B), and Ras proteins (H-Ras and R-Ras). Changes in RASAL3, RASA3, RasGEF1B, H-Ras, and R-Ras mRNA (B) expression and Ras GTPase-activating protein (RasGAP), guanine nucleotide exchange factor (RasGEF), H-Ras, and R-Ras protein (C) expression at the proliferation phase corpus luteum (PPCL, *n* = 4), secretion phase CL (SPCL, *n* = 4), and regression phase CL (RPCL, *n* = 4). mRNA and protein expression in the SPCL and RPCL were normalized to that in the PPCL. **p<0.01 and *p<0.05. The size and molecular weight image of PCR products and proteins were shown as Supplementary Fig S11B and S16B.

### CL morphology, weight, and serum progesterone level during estrous cycle

The PPCL (Fig 5A), SPCL (Fig 5B) and RPCL (Fig 5C) tissues (black arrows) morphology were changed during estrous cycle, and ovulation site of CL diameter (Fig 1D) was higher at SP than PP and RP. The blood vessel (white circle), LEC (white arrows), large LSC (yellow arrows), small LSC (orange arrows) were observed in CL tissue section (Fig 1G, H, I). The number blood vessels were more observed in PPCL (see the supplementary Fig S4A and B) than SPCL (see supplementary Fig S4C, D, E and F) and RPCL (see supplementary Fig S4G and H). The number of large LSC per 10^5^ μm^2^ (Fig 5E) was not be observed in PPCL, whereas could be detected in SPCL and RPCL. Mostly LSCs and LECs were shrunk and damaged in RPCL (Fig 5I) and cell to cell spaces (Fig 5F) were not detected in PPCL and SPCL but could be observed in RPCL. The PPCL was tinged with red (Fig 5J), then size and weight were gradually increased at PP to SP (Fig 5J). Additionally, PPCL inside was redder than SPCL inside, size and weight of RPCL were decreased compared to PPCL and SPCL (Fig 5J and supplementary Fig S5). The RPCL had a yellow color (Fig 5J and supplementary Fig S5C), was harder than PPCL and SPCL (data not shown). The Ras and its regulator factors (Fig 1G, 4B and 4C; Ras activation), tissues size and weight (Fig 5D and 5J; tissue growth), VEGFA, VEGFR2, Ang1 and Tie2 (Fig 3D and 3E; angiogenesis) proteins expression were highest when blood P4 level was consistently increased (Day 2 to 10, PP; see the Fig 5L). On the other hand, Ras activation, tissue growth, and angiogenesis (Fig 5K) were reduced when serum P4 level was highest (Day 14 to 18, SP; see the Fig 5L). However, CL size and weight (Fig 5D and 5J) were rapidly decreased with P4 reduction (Fig 5L), at this point (Day 18 to 20; RP) apoptotic factor proteins (TNFR1, p53, Bax and Casp3; see the Fig 3G and H) were increased compared to PP and SP, most angiogenetic factors and Ras family members were dramatically reduced in RP (Fig 5K).

**Fig 5.**
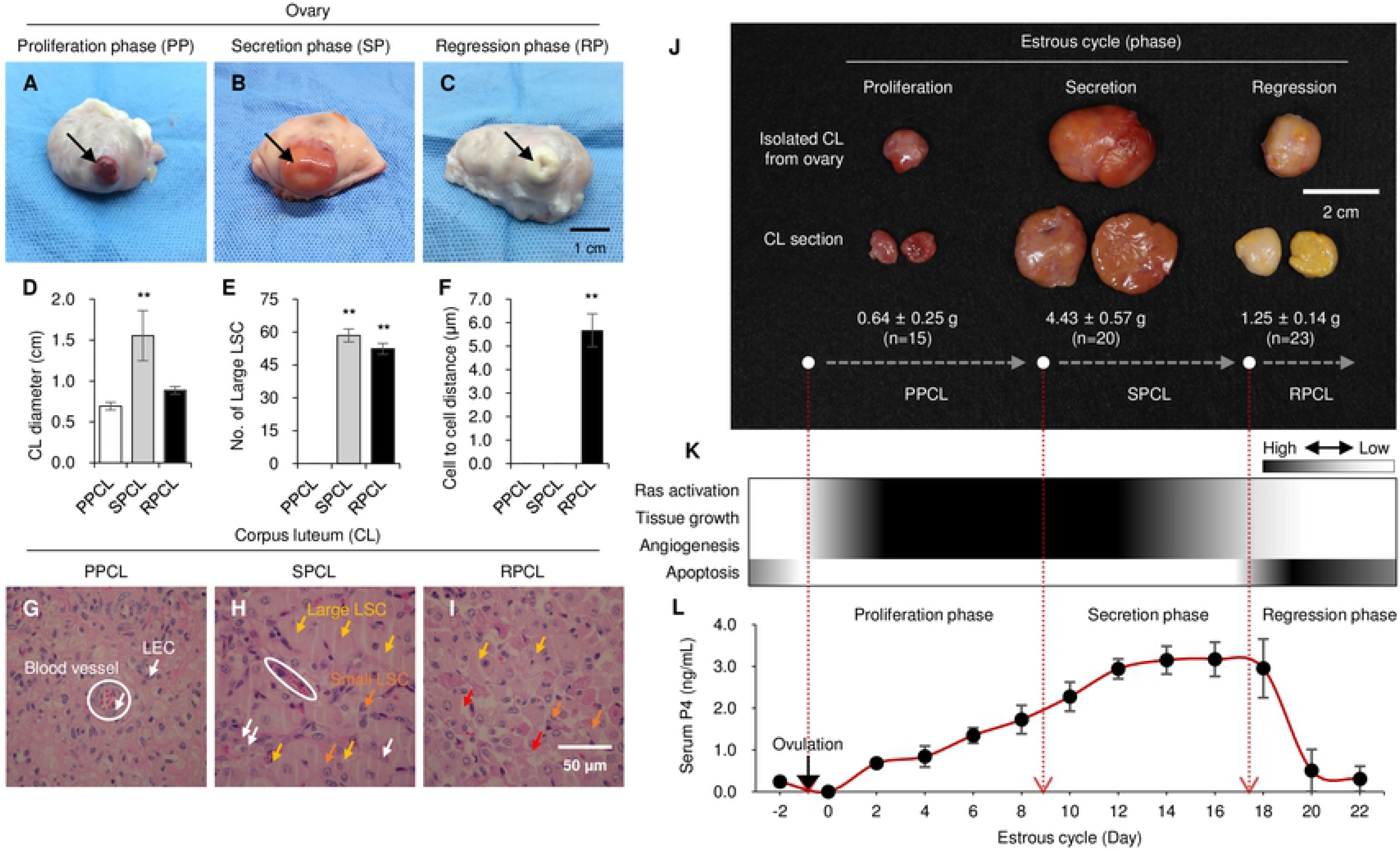
Change of morphology, growth, Ras activation, angiogenesis, apoptosis factors and serum progesterone (P4) level during estrous cycle in corpus luteum (CL). Bovine corpus luteum morphology at the proliferation phase (PP; A), secretion phase (SP; B), and regression phase (RP; C) during the estrous cycle. Black arrows indicate the ovulation site and outside of the corpus luteum (CL; PPCL, *n* = 15; SPCL, *n* = 20; RPCL, *n* = 23) in the ovary; black scale bar = 1.0 cm. Ovulation site of the CL diameter (D), and The PPCL (G), SPCL (H) and RPCL (I) section was evaluated using hematoxylin and eosin staining; blood vessel (white circles), white arrows (luteal endothelial cells; LECs), yellow arrows (large luteal steroidogenic cell; large LSC), orange arrows (small LSC), red arrows (cell-cell space), white scale bar = 50 μm. the number of large LSCs (E) was greater than 20 μm per 1.0 × 10^6^ μm^2^ (PPCL, not detected; SPCL, 58.4 ± 3.0 and RPCL, 52.3 ± 2.5), and the distance of cell to cell (F) (PPCL and SPCL, not detected and RPCL, 5.68 ± 0.70 μm) was calculated under the microscope. Morphology and weight of isolated CL derived from an ovary at PP, SP and RP (J), Ras activation, tissue growth, angiogenesis, and apoptosis activation during the estrous cycle in the CL (K), Changes in the serum P4 level during the estrous cycle in cows. Blood was collected every 2 days for 24 days (*n* = 10 cows). **p<0.01

## Discussion

Generally, P4 increases the endometrium, is involved in pregnancy maintenance and inhibits follicle growth in the female reproductive system (4, 5). In practice, continuous P4 synthesis from LSCs is needed to ensure the interaction between intercellular P4 activity and its nuclear receptor and to active steroidogenic enzymes (11, 21). The main function of the endocrine gland (P4 synthesis and secretion) is to ensure a source of LSCs, but LECs are also important for the development and maintenance of tissues structure in the CL. The CL is composed of LECs (52.7%), large LSCs (3.4%), small LSCs (26.7%), fibrocytes (10.0%), and other cell types (7.5%). Although the numbers of LECs and pericytes are greater than those of the large and small LSCs, the volume density of large LSCs (40.2%) and small LSCs (27.7%) is greater than that of the LECs (13.3%) (22). Generally, LSCs and LECs in the CL develop rapidly after ovulation, during which time activation of angiogenesis is initiated (23). Commonly, VEGF and Ang1 are considered representative angiogenetic growth factors, and a balance of growth factors and their receptors has been reported to contribute to the proliferation of LSCs and LECs during the proliferation phase (8). Therefore, angiogenetic factors and their receptors have special meaning for CL development as a direct result of LSC proliferation, leading to P4 production. Synthesized P4 in LSCs is secreted from the cellular membrane, and the blood P4 level and CL weight consistently increase during the proliferation phase to immediately before the secretion phase. LSC differentiation and proliferation stop at days 12 to 18 after ovulation. Our results showed that the P4 concentration of Korean native cattle at days 12 to 18 after ovulation was 3.0 - 4.0 ng/mL and that the P4 at the secretion phase was higher than that at the proliferation and regression phases. Although the blood P4 level of Korean native cattle heifers was lower than that of previous reports, we confirmed that the pattern of the serum P4 concentration during the estrous cycle was similar to that of previous reports (24, 25). The high PGF2α level and dramatic reduction induce death signal activation in luteal cells, leading to functional and structural luteolysis, such as reduction of P4 production and secretion and luteal tissue and cell degradation (11). Our results detected a large cellular distance, condensation of cellular morphology, and blood vessel degradation in the RPCL. Overall, differentiation of ovarian cells (granulosa and theca cells) to LSCs (large and small LSCs), LSC and LEC proliferation and the resulting angiogenesis, P4 synthesis and secretion of LSCs, and tissue regression related to luteolysis and apoptosis were sequentially matched to the increase, maintenance and reduction of the blood P4 concentration. Therefore, we speculated that the effects of the change in the blood P4 concentration on proliferation, angiogenesis, and apoptosis of luteal cells and tissues might be closely related to Ras family members, because Ras deeply controls cell proliferation, differentiation, morphology, and survival (14).

The story of small G proteins started more than three decades age and was followed by the discovery of the Ras superfamily (26). The major isoforms H-Ras, K-Ras, and N-Ras are highly conserved but show different biological outputs. Additionally, the Ras family includes the Rap, R-Ras, Ral, and Rheb proteins, which play roles mainly as transduction nodes in various signaling pathways (27). Ras family members in the female reproductive tracts have been studied in the endometrium and ovary and typically include activated epidermal growth factor receptor (EGFR) tyrosine kinase, which stimulates mitogen-activated protein kinase/extracellular-signal-regulated kinase (MAPK/ERK) and induces the cell cycle, an increase in the tumor size, and proliferation in various ovarian cancer cells (28, 29). Additionally, Ras is controlled by a molecular switch by cycling between the inactive Ras-GDP and active Ras-GTP conformations, and GDP/GTP binding is regulated by RasGEF and RasGAP(30). As such, Ras has been widely studied in ovarian cancer, but studies of Ras family members during formation and regression of the CL induced by P4 produced from the ovary have not been reported to date.

RASAL3 and RASA3 belong to the RasGAP family, and RasGEF1B belongs to the RasGEF family (30). The RasGAP members are largely divided into four classes [RASA1/p120GAP, neurofibromatosis type 1 (NF1), GAP1^m^ family, and Synaptic GAP (SynGAP)] according to the Src homology (SH), pleckstrin homology (PH) and protein kinase C2 homology domains (31). The first RasGAP to be characterized was p120 RasGAP or Ras p21 protein activator 1 (RASA1) (31, 32). RASA3 is one member of the GAP1^m^ family, and its PH domains bind to phosphatidylinositol (3,4,5)-trisphosphate (PIP_3_) in the cell membrane, which conducts the role of RasGAP. Conversely, RASAL3 is a member of the SynGAP family, but knockdown and knockout studies of this Ras have not been reported (31). RasGAP protein expression was increased in the SPCL when CL formation was completed this result suggested that the RasGAP protein might regulate inhibition of CL formation, because RasGAP induced Ras inactivation as a switch off (30). RasGEF increases the GTP-bound conformation with Ras and contains Sos, RasGRF, and RasGRP. RasGEF1B is a subfamily of RasGEF(33). Our study, during the PP activation of RasGEF was increased when the luteal cells and CL were proliferating and developing and the blood P4 concentration increased until 3.0 ng/mL. However, RasGAP was increased in the PPCL compared to that in the SPCL when luteal cell and CL development stopped and blood P4 was maintained at a high concentration (up to 3.0 ng/mL). All RasGAP and RasGEF protein spots and mRNA and protein expression levels were decreased in the RPCL; during this phase, loss of tissue weight and size, a reduction of the blood P4 concentration, cell disruption, and an increase in the cell to cell distance and apoptotic factors were observed. Therefore, we suggest that the roles of RasGEF and RasGAP are closely related to the formation and finishing of CL development during the estrous cycle. Moreover, the blood P4 level seems to be associated with RasGAP and RasGEF activation. Although an influence of P4 on the effects of Ras, RasGAP, and RasGEF on CL formation and regression has not been reported, this study may contribute to understanding of the roles of the Ras pathway following hormone changes during tissue formation and regression.

Studies of hormones and their receptors have been widely conducted because formation and regression of the CL into the ovary are repeated in response to various hormone reactions, which control the reproductive cycle (4). Based on the bioinformatics analysis, P4R not only reacted to but its transcription was also regulated by ERα, and the interaction between P4R and ERα was closer than those of PGF2αR, ERα, and OTR with P4 in the PPCL. Similar to P4 and E2, the physiological interaction may be more activated in the PP and SP during cellular proliferation and maintenance. However, no association was found between the hormone receptors and the Ras regulators (RASAL3, RASA3, and RasGEF1B). The hormone receptor and Ras regulator aspect of the in silico assay showed that the hormone receptors were not directly regulated by the Ras regulators, we know that there were few study on its receptors and Ras regulators as evidence assay.

We focused on the VEGFA-VEGFR2 and Ang1-Ti2 systems of angiogenetic factors because these systems were well known key factors for CL formation (23). Generally, relationship between sex hormones and angiogenetic systems plays an important role for proliferation and maintenance in the CL lifespan (34). Furthermore, our study showed that angiogenetic receptors (VEGFR2 and Tie2) and RasGEF1B were highly interrelated. Generally, Ras activation starts from receptor tyrosine kinases (RTKs) and G protein-coupled receptors (GPCRs) in the plasma membrane (30, 35). The RTKs are high-affinity cell surface receptors for many polypeptide growth factors, cytokines, and hormones; VEGFR2 and Tie2 (angiopoietin receptor) are included in the RTKs (36). The RTKs and GPCRs combined with growth factors and hormones lead to RasGEF activation and increase the Ras-GTP conformation, which causes cell proliferation, differentiation, and transcription through ERK, MAPK and various cellular pathways (35). We found that RasGEF1B and the angiogenetic receptors bound and reacted to each other and that RasGEF1B was activated by VEGFR2 and Tie2 of the RTK family. However, no interaction was found between RASAL3 and RASA3 of the RasGAP and angiogenetic factors. We suggest that RasGEF is closely related to angiogenesis of the CL during the PP and SP. Thus, increasing the high P4 concentration in the blood may activate RasGEF1B. A study of the relationship between RasGEF1B and P4 is expected in LECs and LSCs.

The prototypic mammalian Ras proteins (H-, K- and N-Ras) share over 90% sequence homology and have similar activity, although H-Ras is a more potent activator of phosphoinositide 3-kinase (PI3K) than K-Ras (37, 38). Deletion of H-Ras results in apoptosis, cell cycle arrest and loss of tumor size in ovarian epithelial cancer cells, whereas H-Ras upregulation increases the reduction of apoptosis but also leads to proliferation through MAPK and ERK signaling in ovarian cancer cells (28, 39, 40). R-Ras is also a small GTPase of the Ras family that regulates cell survival and integrin activity mainly through regulating vascular regeneration in mammalian muscle, intestine, lung, spleen, spinal cord, bone marrow and skeletal muscle (41). We tested the molecular association of H-Ras and R-Ras on interactions with hormone receptors, angiogenetic and apoptotic factors and Ras regulators because we first determined that RASA3 (Ras p21 protein activation) and RASAL3 (Ras protein activator-like 3) directly regulated H-Ras (also called transforming protein p21) in the bovine CL based on our proteomics techniques. Although K-Ras and N-Ras are also transforming protein p21 similar to H-Ras, K-Ras mRNA expression did not differ between the PPCL and SPCL (Supplementary Fig S7), and the protein association between N-Ras and angiogenetic factors were lower than those of H-Ras (Supplementary Fig S8). Second, because R-Ras was deeply related to vascular regeneration, proliferation, and stabilization, we also studied CL formation focus on angiogenetic influences (41). R-Ras only interacted with angiogenetic receptors (VEGFR2 and Tie2), whereas H-Ras had no molecular action with angiogenetic receptors but did interact with VEGFA and p53. There results suggest that H-Ras may have more molecular actions than R-Ras in cellular processes. Additionally, RASAL3 and RASA3 had low correlations with angiogenetic and apoptotic factors, which indicated that downregulation of Ras was not directly associated with angiogenetic and apoptotic factors. Interestingly, no interaction was found between only hormone receptors and Ras regulators, whereas P4R was linked to Ras activation through the ERα–VEGFA–VEGFR2 or Tie2–GasGEF1B and ERα–p53–H-Ras signal pathways. The results signified that P4 could activate Ras via angiogenetic proteins and p53, although few studies have investigated the effects of hormone receptors on Ras in mammals. The observed association pattern of an increase in Ras family member proteins by angiogenetic factors compared to apoptotic factors indicated that Ras might play a regulatory role via angiogenesis and that the hormone receptors that influenced Ras activation might be more involved in the angiogenesis pathway than the apoptosis pathway. Interestingly, molecular signaling from P4R led to the Ras activation signal through angiogenetic factors (transcription of VEGFA), but p53 activation via P4R stimulation could not active overall Ras family members; instead, p53 had molecular actions, such as binding and a transcriptional reaction with H-Ras.

The changes in the RasGAP (RASAL3 and RASA) and RasGEF (RasGEF1B) 2-DE protein spots were similar to the RasGAP and RasGEF immunoblotting results, and H-Ras and R-Ras were strongly expressed in the SPCL. These results show an effect of the abundant RasGEF expression switch on Ras activations in luteal cells of the PPCL, which increases cellular proliferation and differentiation of ovarian cells to steroidogenic cells when CL formation is complete. Conversely, abundant RasGAP restricts the proliferation and differentiation of mature LECs and LSCs in the SPCL. These results indicated that the balance of RasGAP and RasGEF in terms of Ras activation controls the start and finish of CL formation. The blood P4 concentration gradually increased when abundant RasGEF was present and was maintained at high levels when RasGAP was abundant and RasGEF was deficient. Interestingly, the total H-Ras and R-Ras amounts during the PPCL were higher than those during the SPCL which show that numerous Ras proteins may be necessary for successful proliferation and differentiation of luteal cells during the PP. When determining whether Ras family members impact luteal cells, we should note that changes in Ras family members in LECs and LSCs derived from the CL may occur during the estrous cycle because the CL is composed of various cell types. Our results showed strong RasGEF, H-Ras and R-Ras expression in the CL was accompanied by an increase in P4 expression in the blood during the PP. We suggest that the P4 concentration may affect Ras family members and thus proliferation and differentiation of LECs and LSCs. In addition to P4, follicular stimulation hormone (FSH), LH, E2 and PGF2α control the formation and regression of the CL in mammals. Additionally, tissue growth, angiogenesis, and Ras activation coincided with a gradual increase in the blood P4 level in the CL during the PP, and subsequently Ras activation, CL formation, and angiogenesis were matched to a high blood P4 concentration. Although no influence of P4 on Ras, RasGAP and RasGEF was detected in the LECs and LSCs, these results demonstrate that changes in Ras family members follow the serum P4 concentration during tissue formation and regression and protein interactions with hormone receptors, angiogenesis, apoptosis, Ras regulators, and Ras proteins based on the bioinformatics analysis, which is beneficial to fundamental studies of the relationship of sex hormones with Ras during tissue development.

For more than three decades, studies of Ras family members in various mammalian tissues have increased understanding of the biological functions of Ras, especially the effects of mutated Ras on excessive RTK family activation, integrin factor destabilization, and abnormal angiogenesis; ultimately, these events cause abnormal proliferation and survival in tumors and cancers (28, 31, 39, 41). However, few studies have investigated the influence of sex hormones on Ras biological functions, because sex hormones influence particular tissues, such as reproductive tracts, which have sex hormone receptors (42). Unfortunately, studies of the influence of P4 on the CL have mostly focused on domestic animal productivity in livestock for decades. As a result, reproductive science can completely control survival and death of the CL using hormone treatment. On the other hand, living things control the lifespan of special tissues, such as the “corpus luteum”, using the pituitary gland via the reproductive tissue feedback system and steroidogenic hormone regulation, and similarly massive amounts of information are available for tissue formation and regression in living things. Ras family members are one example of this type of information.

## Conclusions

We suggest that the detailed molecular function of Ras is available for discovery during formation and regression of the CL. Therefore, understanding the roles of the RasGAP, RasGEF, and Ras proteins following shifts in the P4 concentration may provide new perspectives for the relationship between Ras and hormones during tissue formation and regression. We suggest that this knowledge may contribute to therapy for hormonal diseases.

## Acknowledgements

The authors thank the Institute of Animal Resources at the Kangwon National University for their services. This work was supported by National Research Foundation of Korea (NRF 2018R1D1A3B07048167), Republic of Korea. This study supported by 2017 Research Grant from Kangwon National University (No. 520170021).

## Author contribution

S.H. Lee and S. Lee designed and oversaw the study. S.H. Lee collected samples, S.H. Lee and S. Lee performed experiment. S. Lee analyzed genes and proteins, and S.H. Lee performed proteomics and bioinformatics. S.H. Lee wrote the paper and S. Lee made figures and tables. S.H. Lee and S. Lee wrote figure legends and supplementary information. All authors contributed to the critical review of the manuscript.

## Abbreviations

2-DE: two-dimensional electrophoresis
CL: corpus luteum
P4: progesterone
PP: proliferation phase
RP: regression phase
SP: secretion phase

